# An N4-like *Caulobacter* phage requires host smooth lipopolysaccharide biosynthesis for infection

**DOI:** 10.1101/2025.10.24.684376

**Authors:** Maeve McLaughlin, Katheren Barger, Charlotte Barron, Makena Fisher, Priscilla Mac-Kittah, Aretha Fiebig, Sean Crosson

## Abstract

*Caulobacter* species are Alphaproteobacteria that commonly inhabit plant-associated and aquatic microbial communities. Although *Caulobacter* is widespread and has long served as a model for the study of bacterial cell biology, our understanding of the diversity of viruses that infect *Caulobacter* species is limited. Here, we describe the discovery and characterization of Circe, a freshwater N4-like podophage belonging to the *Schitoviridae* family that infects *C. crescentus*. We isolated two variants, CirceC and CirceH, that differ by a single nucleotide resulting in an F91I substitution in Gp063, an uncharacterized protein found in diverse bacteriophages and bacteria. While both Circe variants adsorb to *C. crescentus* with similar efficiency, they produce morphologically distinct plaques and display different infection dynamics. Through forward genetic selection and genome-wide transposon fitness profiling, we identified *C. crescentus* genes involved in cell envelope assembly, membrane sphingolipid biosynthesis, and envelope polysaccharide biosynthesis that influence susceptibility to Circe infection. Loss-of-function mutations in a predicted nucleoside diphosphate sugar epimerase and multiple genes required for smooth lipopolysaccharide (S-LPS) biosynthesis and export conferred strong resistance to infection. These results support a model in which S-LPS functions as a receptor for phage Circe. Our study expands the known repertoire of *Caulobacter* phages and adds to a growing understanding of the role of envelope polysaccharides in bacterial infection by N4-family phages.

**IMPORTANCE:** Viruses that infect bacteria and archaea, known as phages, shape microbial community structure and function. Phages initiate infection by binding to specific molecules on the surface of host cells. Yet for many microbes, the identities of infecting phages and their corresponding host receptors remain poorly defined. *Caulobacter* spp. are ecologically important bacteria that can produce a variety of protein surface structures, including pili, a flagellum, and a surface layer (S-layer), all of which have been identified as phage receptors in this genus. We discovered *Caulobacter* phage Circe and provide evidence that it relies on host smooth lipopolysaccharide to infect *C. crescentus*. This study broadens understanding of phage-host interactions in *Caulobacter* and establishes Circe as a new system to investigate the molecular mechanisms by which phages engage with bacterial cells.

## Introduction

*Caulobacter* is a widely distributed genus of Alphaproteobacteria found in diverse environments, including aquatic habitats, soils, and plant-associated niches (1, 2). Long studied as a model system for bacterial cell biology (3), *Caulobacter* spp. are now also recognized as important members of plant-associated microbiomes (4) with select strains shown to promote plant growth (5, 6). The diverse ecological roles of *Caulobacter*, together with its genetic tractability, make it a powerful system for studying how environmental interactions impact bacterial biology.

Understanding how *Caulobacter* interacts with its environment requires consideration of the biotic forces that shape its physiology, ecology and evolution. Among the most significant of these are bacteriophages, which are viruses that infect bacteria. With an estimated global abundance of approximately 10^31^ particles, bacteriophages (or phage) are the most numerous biological entities on Earth (7). Phages play critical roles in microbial communities by regulating host population sizes, driving nutrient turnover, and mediating genetic exchange in communities (8-11). Infection of a host cell by a phage typically begins with highly specific binding to a surface receptor molecule such as a protein or polysaccharide. These receptor-mediated interactions exert intense selective pressure on bacterial populations and frequently lead to the emergence of phage-resistant mutants with altered cell surface properties (12-14). Several phages that infect *Caulobacter* are known. The virulent siphophage, φCbK, was among the first *Caulobacter* phages reported (15) and has since been used extensively to study polar morphogenesis due to its dependence on the polar flagellum and pili for infection (16). The T4-like phage, φCr30, uses the paracrystalline surface-layer protein as a receptor (17) and remains a useful tool for transduction of *Caulobacter* (18). Other double-stranded DNA and RNA *Caulobacter* phages have also been described (19-21), but the broader diversity and ecological roles of *Caulobacter* phages remain poorly defined.

The N4-like group of phages represents a distinct viral lineage that was first isolated on *Escherichia coli* in 1966 (22). Phage N4 is a podovirus characterized by an icosahedral capsid, a short non-contractile tail, and a linear double-stranded DNA genome of approximately 70 kb (23, 24). A defining feature of this group is a large virion-associated RNA polymerase (vRNAP), which is co-injected with the genome to initiate early transcription in the host cell (25-28). N4 coliphage are also notable for their delayed lysis strategy, which enables exceptionally high burst sizes, often exceeding 3,000 plaque-forming units (PFU) per infected cell (29).

Here, we report the discovery and characterization of Circe, an N4-like phage of the *Schitoviridae* family that infects *C. crescentus*. To our knowledge, Circe is the first N4-like phage described to infect a *Caulobacter* species. We isolated two phage variants, CirceC and CirceH, that differ by only a single nucleotide yet have distinct plaque phenotypes and infection kinetics in a broth infection assay. Through a combination of genetic selections and screening approaches, we demonstrate that successful infection by Circe requires several genes that function in cell envelope carbohydrate metabolism. Specifically, we show that a WcaG-like NAD-dependent sugar epimerase and multiple genes required for LPS O-polysaccharide biosynthesis as essential host factors for Circe infection. These results expand our understanding of N4-like phage biology, reveal new molecular determinants of phage susceptibility in *C. crescentus*, and establish Circe as a tractable model for studying phage-host interactions in an ecologically important genus.

## Results

### Isolation and structural characterization of *Caulobacter* phages

While measuring *Caulobacter crescentus* CB15 growth in filtered water collected from an inland freshwater lake in Haslett, Michigan (USA), we unexpectedly observed zones of clearing on spot plates used for colony-forming unit (CFU) enumeration. These clearing zones were consistent with lytic bacteriophage activity and were sampled for further analysis. Subsequent experiments confirmed that the clearing was caused by phage capable of infecting and lysing *C. crescentus* CB15 on solid medium.

Two distinct plaque morphologies were observed: (1) clear plaques and (2) halo plaques characterized by a clear center and turbid outer edge (Figure 1A). Phage isolated from each plaque type consistently reproduced only the original plaque morphology, suggesting that the clear and halo plaques were generated by distinct phages. We purified and imaged phage particles from each plaque type using transmission electron microscopy (TEM). The phage from the clear plaques, and the phage from the halo plaques both displayed capsids with mean diameters of 79.8 ± 5.4 nm (n=18) and 69.5 ± 3.5 nm (n=17), respectively (Figure 1B–C). Neither phage exhibited a prominent tail structure.

**Figure 1.**
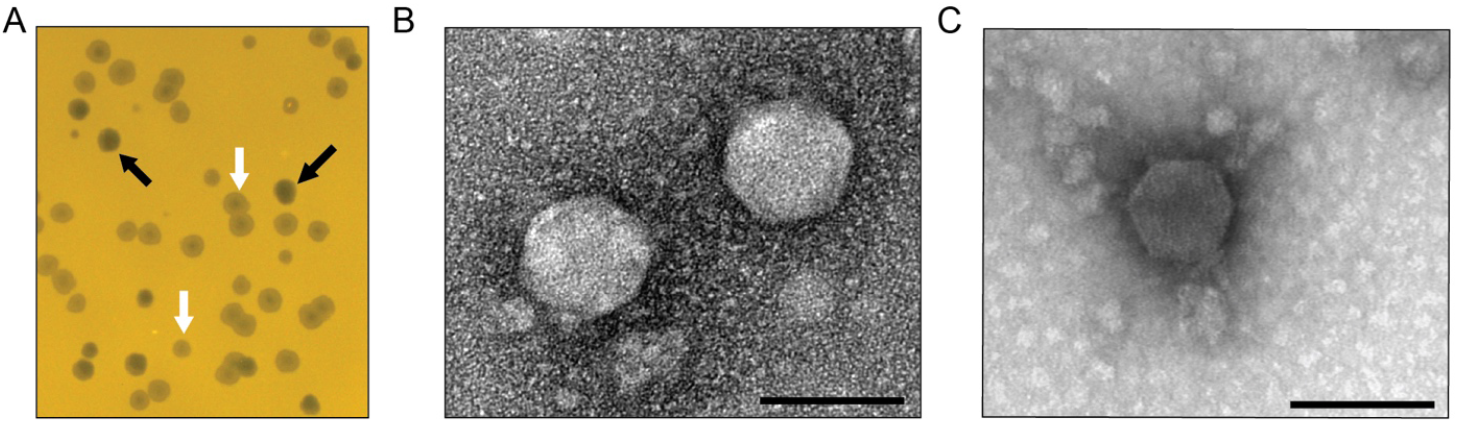
Isolation and structural characterization of *Caulobacter*-infecting phages from a freshwater lake. (A) Representative plaque morphologies formed by Lake Lansing phages on *Caulobacter crescentus* CB15 grown on PYE agar medium. Clear plaques (black arrows) and halo plaques with turbid outer rings (white arrows) were consistently observed, suggesting two distinct phage types. (B–C) Transmission electron micrographs of negatively stained phage particles purified from clear plaques (B) and halo plaques (C), both displaying podovirus morphology characterized by icosahedral capsids and short tails. Scale bars: 100 nm.

### Genomic features of clear-and halo plaque-forming variants of *Caulobacter* phage Circe

We isolated and purified phage from clear and halo plaques and sequenced their genomes. Both phage genomes comprised a 73,793 bp linear double-stranded DNA genome with 341 bp direct terminal repeats and a G+C content of 53%, which is notably lower than that of the *C. crescentus* host genome (67.2%; GenBank accession CP001340). There was only a single base pair difference between the clear plaque- and halo plaque-forming phage: an A-to-T transversion at position 38,705 in the isolate that forms halo plaques. Genome annotation (see Materials and Methods) predicted 100 open reading frames (ORFs) and 2 tRNA genes (Table S1). Accordingly, we designate the halo plaque-forming isolate *Caulobacter* phage CirceH (hereafter, CirceH) and the clear plaque-forming isolate *Caulobacter* phage CirceC (hereafter, CirceC).

Phylogenetic analysis (15) placed Circe within the *Schitoviridae* family, clustering it with N4-like phages. The N4 phages are defined by the presence of seven hallmark genes (24): a DNA polymerase, a large virion-associated RNA polymerase (vRNAP), a portal protein, a major capsid protein, a tail protein, and both the large and small terminase subunits; Circe contains all seven hallmark genes. Analysis of genes associated with host cell lysis revealed four ORFs with predicted roles in this process. Among these was *gp077*, predicted to encode an Rz-like inner membrane spanin (i-spanin). An outer membrane spanin (o-spanin) was not initially evident in the genome, but a *tblastn* search using the o-spanin from *Escherichia* phage N4 identified a candidate gene in the +1 reading frame relative to *gp077*. This gene, designated *gp078*, shared 52% overall identity with the periplasmic domain of the N4 o-spanin and possessed a predicted outer membrane lipoprotein signal sequence, supporting its identity as a Circe o-spanin. Downstream of the spanin system, *gp079* was annotated as a glycoside hydrolase family endolysin (IPR002196), likely i nvolved in peptidoglycan degradation. While no holin or antiholin genes were definitively identified, three downstream ORFs (*gp080–gp082*) encoded predicted transmembrane domains, raising the possibility that they function in membrane disruption during lysis. As noted above, the only genomic difference between CirceH and CirceC is a single A-to-T transversion at position 38,705 in CirceH, resulting in a phenylalanine-to-isoleucine (F91I) substitution in *gp063*, a gene of unknown function. The mechanism by which this mutation may impact plaque morphology is discussed in a later section.

### Host genetic background and phage genetic variation impact infection phenotypes

To evaluate the infection dynamics of the two Circe variants in closely related *C. crescentus* strains with distinct cell envelope features, we measured the efficiency of plating (EOP) of CirceC (clear plaque-forming) and CirceH (halo plaque-forming) on strain NA1000. NA1000 is a laboratory strain that is nearly identical to CB15; the two strains descend from a common ancestral isolate (30) and differ by only eight single-nucleotide polymorphisms (SNPs), two single-base insertions or deletions, and the presence of a 26-kb mobile genetic element (MGE). Compared to the *C. crescentus* CB15 strain, both CirceC and CirceH formed fewer and more turbid plaques on NA1000 (Figure 3), suggesting reduced infectivity. Previous work has shown that the MGE in strain NA1000 encodes a suite of enzymes involved in exopolysaccharide (EPS) biosynthesis, and that this EPS production enhances resistance to the S-layer-targeting phage φCr30 (30, 31). To test whether the MGE also affects susceptibility to CirceC and CirceH infection, we examined EOP on an NA1000 derivative lacking the MGE (ΔMGE) (30). On the NA1000 ΔMGE strain, both phages formed plaques comparable in number and clarity to those observed on CB15, indicating that the loss of MGE-encoded EPS biosynthesis restores phage susceptibility and that the MGE contributes to resistance aga inst Circe infection.

**Figure 2.**
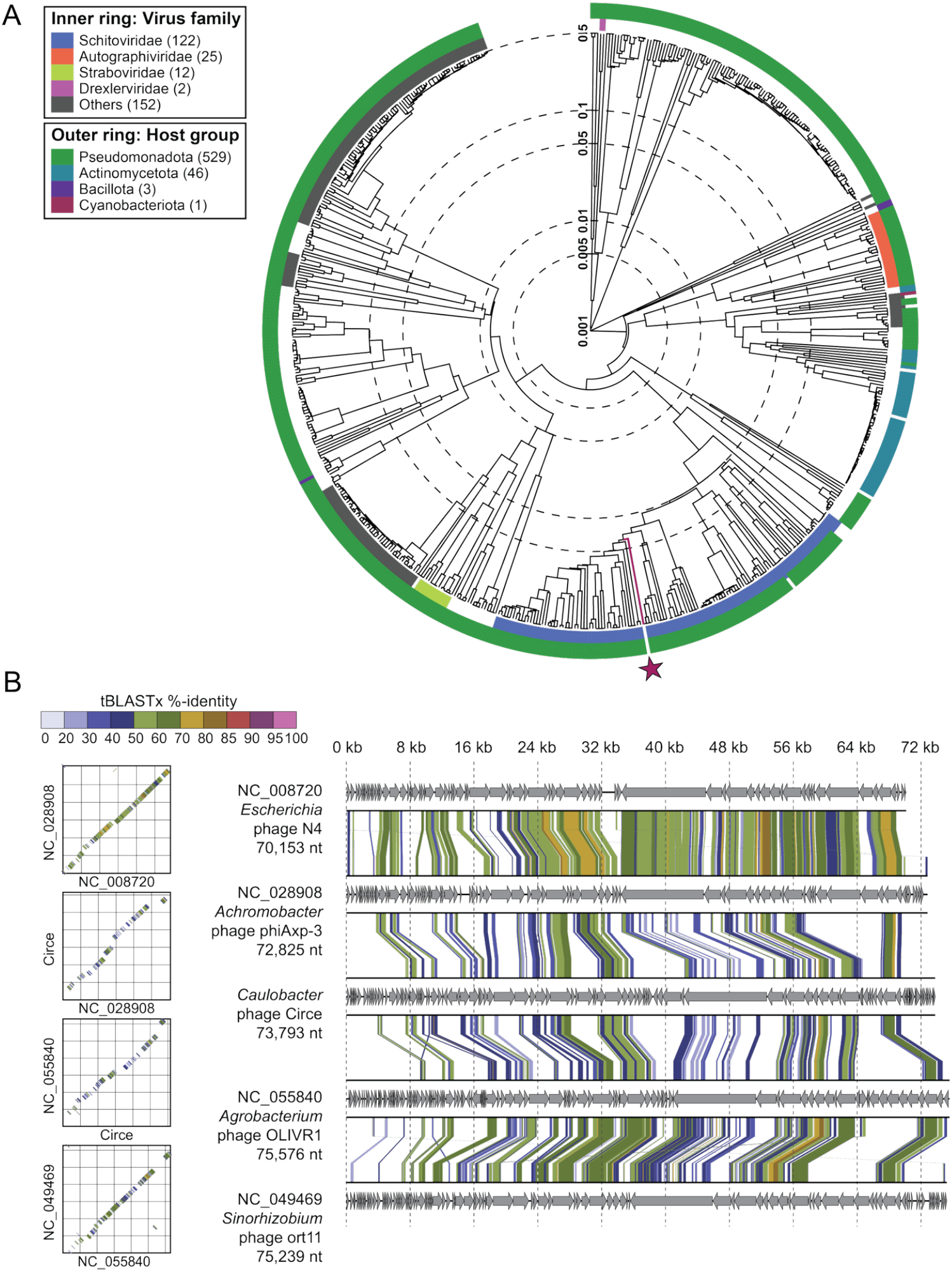
Circe is a member of the N4-like Schitoviridae. A) Viral proteomic tree generated using ViPTree (http://www.genome.jp/viptree), showing the relationship between Circe and 590 related phage genomes. The inner ring denotes viral family-level classification, while the outer ring indicates host taxonomic group. Circe is highlighted with a red line and red star. B) Genome alignments and dot plots comparing Circe to representative N4-like phages. Arrows represent predicted genes in each genome, and connecting lines denote regions of nucleotide similarity based on tBLASTx comparisons. Color intensity corresponds to percent identity, illustrating conserved synteny across the N4-like lineage. (note: *Caulobacter* phage Circe is awaiting an NCBI genome accession number).

**Figure 3.**
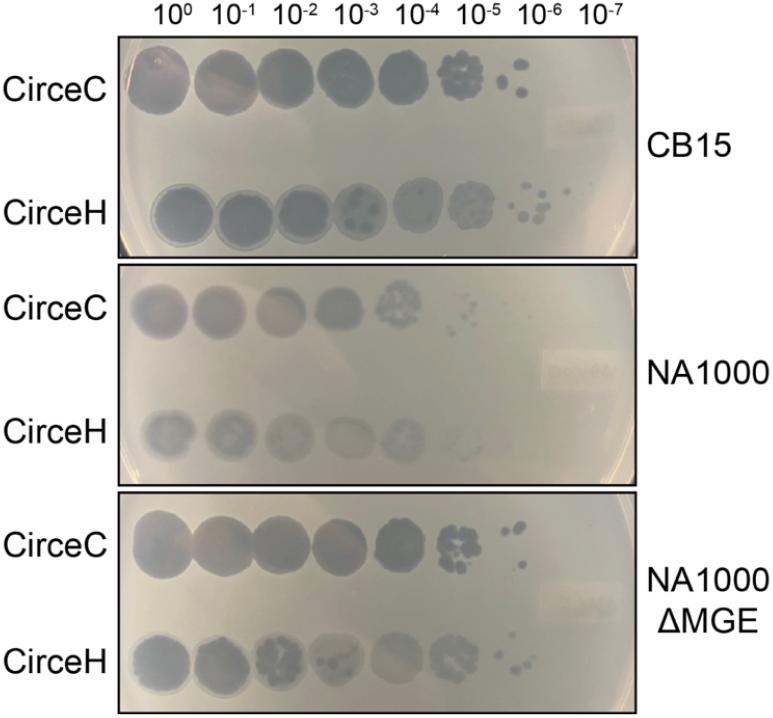
A mobile genetic element (MGE) in *C. crescentus* NA1000 reduces susceptibility to phage Circe. Serial 10-fold dilutions of CirceC (clear plaque-forming) and CirceH (halo plaque-forming) were spotted on PYE top agar containing Caulobacter crescentus strains CB15 (top panel), NA1000 (middle panel), and an NA1000 derivative that lacks a 26 kb mobile genetic element (NA1000 ΔMGE, bottom panel). Circe phages formed fewer and more turbid plaques on NA1000 relative to CB15, but plaque formation was restored in the ΔMGE strain, suggesting that the MGE contributes to phage resistance. Image is representative of least three biological replicates.

We next compared the ability of CirceC and CirceH to infect and lyse *C. crescentus* in liquid culture. Exponentially growing CB15 cells were infected at multiplicities of infection (MOIs) of 0.1, 1, and 10, and optical density at 660 nm was tracked over time (Figure 4). Infections with CirceC produced a characteristic delay in growth suppression followed by a rapid decline in absorbance, consistent with lytic activity. The timing of lysis was MOI-dependent: lysis began approximately 5 hours post-infection at MOI 0.1, 3 hours at MOI 1, and within 1 hour at MOI 10. In contrast, CirceH infections followed a more complex pattern. While lysis occurred at all MOIs, the decline in optical density was slower and biphasic. At MOI 10, absorbance initially decreased from 1 to 3 hours, then partially rebounded from 3 to 6.5 hours before declining again and fully clearing the culture by 10 hours (Figure 4). Similar delayed or multiphasic trends were observed at lower MOIs. These results suggest that despite near-identical genomes, CirceH exhibits altered infection dyna mics compared to CirceC, possibly reflecting differences in adsorption efficiency, replication kinetics, or inter actions with host cell surface structures.

**Figure 4.**
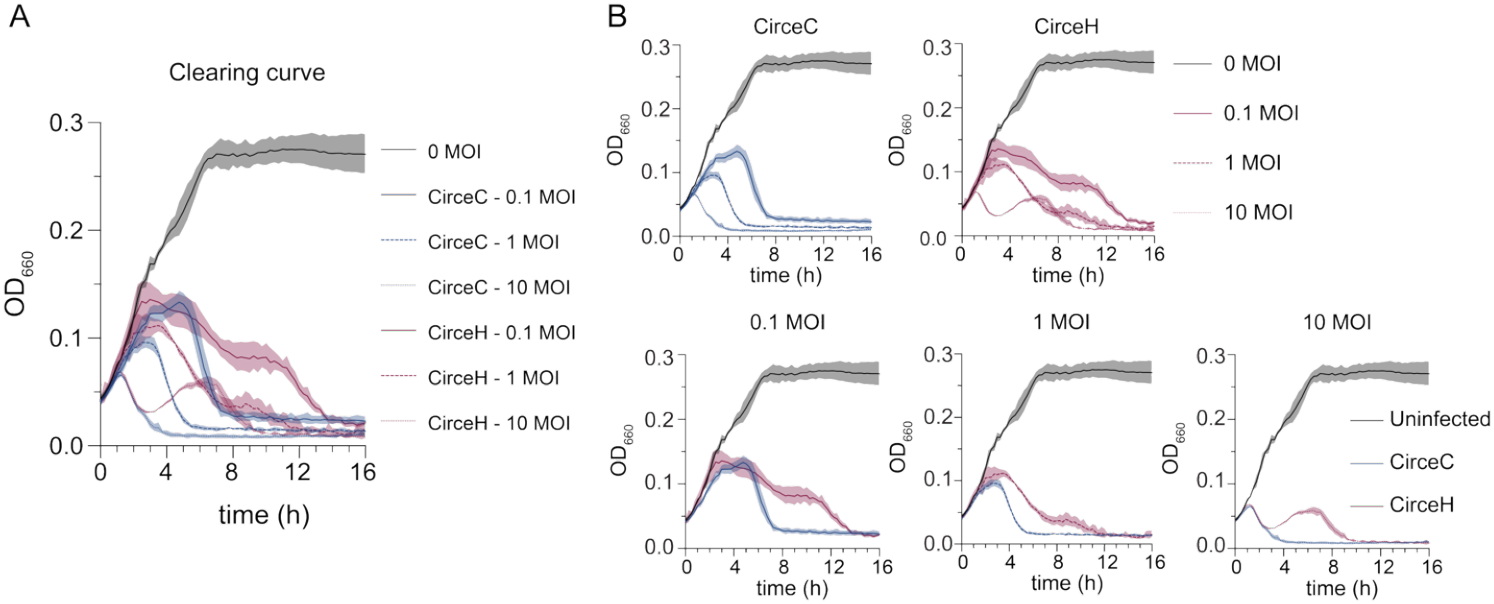
CirceC and CirceH exhibit distinct clearing kinetics during infection of *C. crescentus*. A) *C. crescentus* CB15 broth cultures were infected with either CirceC or CirceH at multiplicities of infection (MOIs) of 0.1, 1, or 10 in complex PYE medium. Optical density at 660 nm (OD_660_) was measured every 15 minutes for 16 hours to track lysis dynamics. Uninfected cultures served as controls. CirceC triggered a sharp, MOI-dependent drop in OD_660_ consistent with rapid lysis, while CirceH exhibited slower, biphasic clearing kinetics. B) The data from panel A are reorganized by phage (top row: CirceC, CirceH) and MOI (bottom row: 0.1, 1, 10) to highlight kinetic differences. Lines represent the mean, and shaded areas represent standard deviation from three biological replicates.

### CirceC and CirceH have equivalent host adsorption properties

We postulated that the differences in infection dynamics between the two Circe variants were due to differences in their ability to adsorb to host cells. We therefore quantified adsorption of CirceC and CirceH to *C. crescentus* CB15 over a 60-minute time course by measuring the fraction of unadsorbed phage remaining in culture supernatants. Both variants displayed similar adsorption properties (Figure 5). For CirceC, approximately 87 % of phage particles were adsorbed within 30 minutes of incubation, and adsorption reached near completion by 60 minutes, yielding an adsorption rate constant of k = 4.2 × 10^−10^ mlmin^−1^. CirceH exhibited a comparable profile, with 86 % adsorption by 30 minutes and a calculated rate constant of k = 4.3 × 10^−10^ mlmin^−1^. We conclude that the F98I substitution in *gp063* distinguishing CirceH from CirceC does not measurably alter the efficiency or kinetics of phage adsorption to *C. crescentus* CB15.

**Figure 5.**
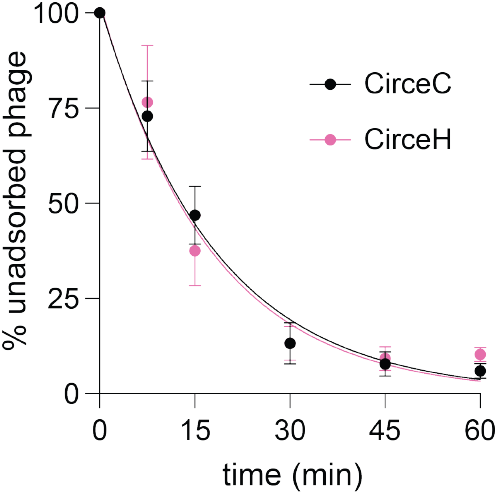
Adsorption kinetics of CirceC and CirceH to *C. crescentus* CB15. **(A)** Exponentially growing *C. crescentus* CB15 cultures were infected with either CirceC (black circles) or CirceH (pink circles) at a multiplicity of infection (MOI) of 0.01 in PYE medium. At the indicated time points post-infection, the fraction of unadsorbed phage in the supernatant was quantified and expressed as a percentage relative to the initial phage titer at time zero. Data represent the mean ± standard deviation of three biological replicates. Curves represent fits to the model: P(t) = P_0_ × exp(–k × n × t), where P(t) is the concentration of unadsorbed phage at time t, P_0_ is the initial phage concentration, n is the bacterial cell density, t is time (minutes), and k is the adsorption rate constant. The R^2^ goodness of fit for each regression: CirceH =0.94; CirceC =0.97.

### Loss-of-function mutations in a WcaG-like epimerase gene confer resistance to Circe

In the process of conducting plaque assays with CirceC, we observed that *C. crescentus* colonies occasionally emerged within lysis zones after two days of incubation. To test whether these colonies were resistant to CirceC, we isolated three representative clones and measured their susceptibility to infection. All three mutants exhibited complete resistance to CirceC: no plaques were observed at any phage concentration (Figure 6A), and cultures inoculated with 0.1 MOI of CirceC grew comparably to uninfected controls in liquid culture (Figure 6B). To identify the genetic basis for resistance, we sequenced the genomes of these three spontaneous mutants and compared them to the parental *C. crescentus* CB15 strain. Each mutant harbored a distinct mutation in a gene encoding a WcaG-like NAD-dependent epimerase (gene locus tag *CCNA_03538* or *CC_3425*) (Table S2). Hereafter, we refer to *C. crescentus* gene locus tags using the CCNA_ prefix, which corresponds to our preferred *C. crescentus* genome annotation.

**Figure 6.**
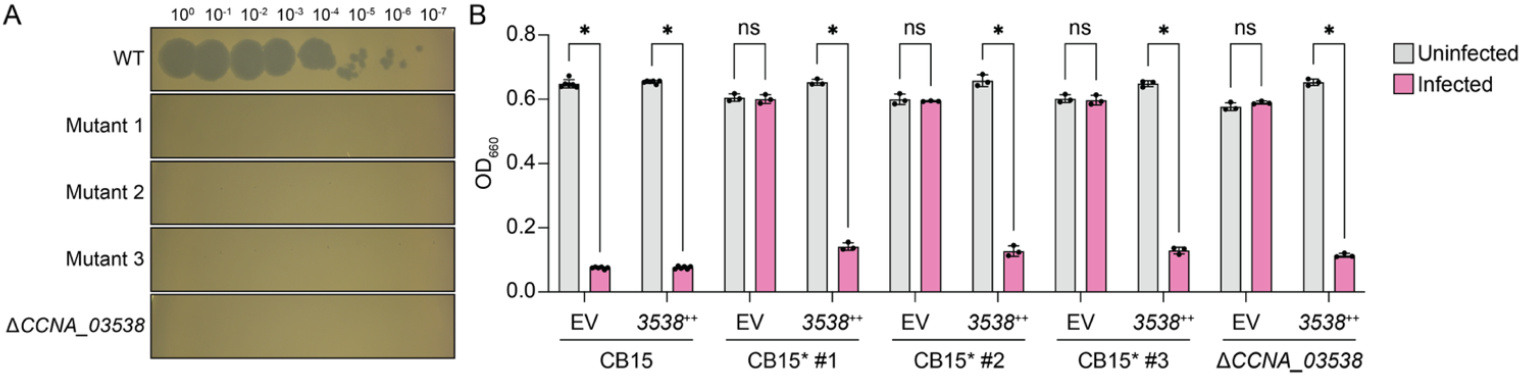
Mutation or deletion of a putative WcaG-family epimerase (CCNA_03538) confers resistance to Circe infection. (A) Plaque assay showing CirceC infection of wild-type *Caulobacter crescentus* CB15, three independently-isolated spontaneous Circe-resistant *C. crescentus* mutants (Mutant 1–3), and a *C. crescentus* strain with an in-frame deletion of the WcaG-family sugar epimerase gene *CCNA_03538* (Δ*CCNA_03538*). Tenfold serial dilutions of CirceC were spotted onto PYE top agar seeded with the indicated strains. (B) Growth assay in liquid culture following infection with CirceC at MOI 0.1. Optical density at 660 nm (OD_660_) was measured after 24 hours of incubation. Each strain contained either an empty vector (EV) or a plasmid expressing *CCNA_03538* under an inducible promoter (++). Expression of *CCNA_03538* restored phage sensitivity in all resistant strains. Bars represent the mean ± SD from three biological replicates. Statistical significance was assessed using a two-way ANOVA with post test (ns, not significant; *p* < 0.05).

To test whether *CCNA_03538* is necessary for CirceC infection, we generated an in-frame deletion strain (Δ*CCNA_03538*) and assessed its phage sensitivity. Like the spontaneous mutants, Δ*CCNA_03538* was completely resistant to CirceC infection in both plaque and liquid assays (Figure 6A–B). Genetic complementation of either the inframe deletion strain or the spontaneous mutants with *CCNA_03538* expressed from a cumate-inducible plasmid (32) restored phage susceptibility, confirming that loss of *CCNA_03538* function confers CirceC resistance (Figure 6B).

### Genome-Wide Identification of Circe Host Factors by RB-TnSeq

To systematically identify additional host genes required for Circe infection, we performed a phage challenge experiment using a randomly barcoded transposon (RB-TnSeq) library of *C. crescentus* CB15. The library was infected with CirceC at a multiplicity of infection (MOI) of 1, and barcode abundance was quantified at multiple time points across the infection cycle (Figure 7A). As expected, transposon insertions in the *CCNA_03538* epimerase were highly enriched by 7.5 hours post-infection, consistent with the strong resistance phenotype observed in both spontaneous mutants and in-frame deletions (Figure 7B, Table S3). In addition, we observed late enrichment of transposon mutants in multiple genes required for LPS O-polysaccharide biosynthesis, including *CCNA_00497, CCNA_01068, CCNA_02386, CCNA_03733*, and *CCNA_03744*; these genes have been previously implicated in assembly or export of smooth lipopolysaccharide (S-LPS) (33, 34) (Figure 7; Table S3). Several of these mutants also exhibited mild fitness defects at early time points, potentially reflecting increased vulnerability to phage infection stress (in cases where mutants become infected). This result may also simply reflect slower growth of these mutants at earlier time points. By late time points (7.5 h), however, the large competitive advantage of S-LPS and other envelope polysaccharide mutants is evident. To validate screen results showing that S-LPS genes are important host factors for Circe infection, we constructed inframe deletions in *CCNA_00497, CCNA_02386*, and *rfbB* and tested their susceptibility to CirceC. Each mutant was resistant to infection in liquid-based assays (Figure 7C). Genetic complementation with each corresponding gene from a xylose-inducible promoter restored phage sensitivity, confirming smooth LPS is a key host contributor to Circe infection.

**Figure 7.**
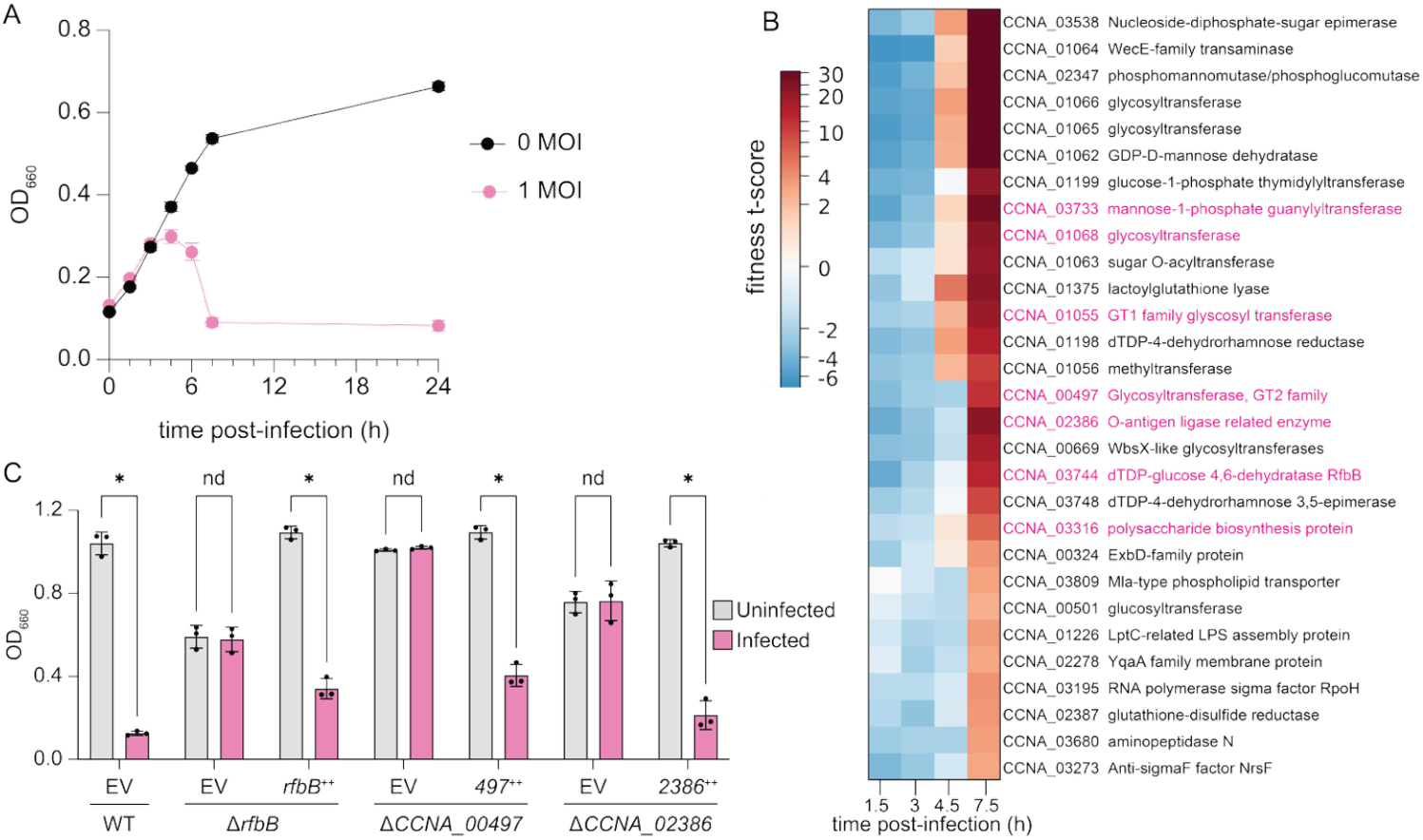
Disruption of smooth LPS and other cell envelope polysaccharide genes confers resistance to Circe infection. A) Growth of a *C. crescentus* CB15 randomly barcoded transposon (RB-TnSeq) library in the presence or absence of Circe. Cultures were infected at an MOI of 1 during exponential phase in PYE medium, and optical density (OD_660_) was monitored over 24 hours. B) Excerpt of a hierarchically clustered heatmap of gene fitness t-scores at 1.5, 3.0, 4.5, and 7.5 h post-infection. Panel shows the cluster containing genes (highlighted in pink) with known associations with smooth lipopolysaccharide (S-LPS) biosynthesis. Positive scores indicate mutants with relatively enhanced fitness at a given time point; mutants with negative scores had relatively reduced fitness. Complete fitness data are presented in Table S3. C) Validation of Circe resistance in in-frame deletion mutants of *rfbB, CCNA_00497*, and *CCNA_02386*. Strains were infected at MOI 0.1 in liquid PYE cultures and OD_660_ was measured after 24 hours. Each strain carried either an empty vector (EV) or a plasmid expressing the deleted gene (++). Bars represent the mean ± SD from three biological replicates. Statistical comparisons between uninfected and infected conditions were made using an unpaired two-tailed t-test (p < 0.05; ns = not significant).

We further identified a distinct group of genes that showed fitness advantages at early time points post-infection that decreased by 7.5 hours, suggestive of host functions that impact the early stages of infection or that are more relevant at low phage titer (Figure S1). This class included genes from the sphingolipid biosynthesis and export cluster spanning *CCNA_01210* to *CCNA_01226* (35-37). Specific mutants in this cluster including *CCNA_01210* (phosphoglycerate cytidylyltransferase), *CCNA_01212* (ceramide synthase), and *CCNA_01220* (serine palmitoyltransferase) showed an interesting fitness profile with modest but reproducible fitness gains that peaked at 4.5 hours but that returned to near zero by 7.5 hours. These data provide evidence that membrane sphingolipids contribute to efficient CirceC infection. It is known that *C. crescentus* lipid A is dispensable under select genetic conditions, if sphingolipids are present (34). The relationship between the *C. crescentus* sphingolipid biosynthesis gene cluster, LPS O-polysaccharide synthesis, and susceptibility to phage Circe is an interesting future area of investigation.

Our genome-scale screen also revealed a group of genes that exhibited basally reduced fitness scores across all time points, including the 0 h time point (Table S3), likely reflecting a general growth defect. These genes included CCNA_01427 (*bamE*), a lipoprotein subunit of the β-barrel assembly machinery (BAM complex) previously shown to have basally low fitness scores in Tn-seq studies [5]; cell envelope regulator CCNA_01817 (*ntrX*) (38); CCNA_00050 (*lnt*), which catalyzes the final step in outer membrane lipoprotein maturation, and CCNA_02037 (*lon*), encoding an ATP-dependent protease involved in general proteostasis.

### Discussion

### On the significance of sequence variation in Circe gene gp063

Circe is a member of the *Schitoviridae* family and, to our knowledge, represents the first N4-like phage reported to infect *Caulobacter*. We isolated two Circe variants (CirceC and CirceH) that differ by a single nonsynonymous mutation in the gene gp063. A PSI-BLAST search identified homologs of gp063 in several *Caulobacter* and *Bradyrhizobium* genomes, as well as in the temperate *Caulobacter* phage S2B. In some of these genomes, Gp063 homologs have been generically annotated as tail fiber proteins. However, we have found no experimental evidence that Gp063 or its homologs contribute to tail fiber structure or function in any phage. To further explore the possible function of gp063, we generated a predicted structure using AlphaFold3 (39) (Figure S2A) and queried it against the BFVD database (40) using FoldSeek (41). This analysis identified structural similarity to several Caudovirales phage proteins, which share low sequence identity (∼20%) with gp063 but exhibit highly similar predicted tertiary structures. gp063 did not share significant structural homology with any proteins of known function in other structural databases accessed by Foldseek.

Although CirceC and CirceH adsorbed to *C. crescentus* with similar efficiency, CirceH displayed delayed and biphasic lysis kinetics, suggesting that the Gp063 mutation affects a post-adsorption stage of the infection cycle (e.g. injection, replication timing, or lysis regulation). Our structural predictions indicate that the C-terminal domain of Gp063 contains five α-helices, with five phenylalanine residues buried within a hydrophobic core (Figure S2B). In CirceH, one of these stacked phenylalanines (F91) is replaced by an isoleucine (Figure S2B). This substitution may disrupt the packing of the α-helical bundle, potentially altering Gp063 function and contributing to the observed phenotypic differences in plaquing and in broth culture infections. Although it remains unclear which Circe variant represents the ancestral or predominant form in nature, the presence of two *gp063* alleles that yield phage with distinct infection phenotypes suggests potential for adaptive diversification at this locus. The coexistence of CirceC and CirceH aligns with established phage strategies for modulating host range, such as phase variation at tail fiber loci that alters expression or structural properties (42, 43), or the use of diversity-generating retroelements to modify receptor-binding proteins (44). It is plausible that *Caulobacter* phage Circe populations experience selection on *gp063* variants that enable adaptation to changes in local receptor landscapes or that help maintain fitness in the face of other host population shifts.

### Circe employs a host engagement strategy characteristic of other N4-like phages

To identify host factors required for infection by *Caulobacter* phage CirceC, we used forward genetic selection and genome-wide RB-TnSeq screening. These approaches identified several host genes critical for productive infection. Among the strongest hits were genes involved carbohydrate metabolism, including genes necessary for smooth lipopolysaccharide (S-LPS) biosynthesis and export. Mutations in several of these genes conferred complete resistance to Circe, supporting a model in which one or more components of S-LPS act as the primary adsorption receptor. This result aligns with established mechanisms of host engagement by other N4-like phages. For example, *Escherichia* phage N4 initially binds an exopolysaccharide produced by NfrB before transitioning to NfrA, an outer membrane protein, as its terminal receptor (45-48). Similarly, lipopolysaccharide has been shown to mediate host interaction for N4-like phages infecting *Achromobacter* and *Shigella* species (49, 50). Our results now extend this model to *Caulobacter*, indicating that cell envelope polysaccharide recognition may be a conserved feature across diverse N4-like phages.

In *C. crescentus*, S-LPS also anchors the paracrystalline surface layer (S-layer) protein RsaA (51), which forms a highly ordered envelope surface structure (52). Although prior studies have proposed that the S-layer can shield phage receptors and inhibit infection, our RB-TnSeq screen did not detect a fitness advantage for *rsaA* mutants during Circe challenge (Table S3). This indicates that Circe does not require the S-layer for adsorption and likely engages S-LPS directly. This requirement distinguishes Circe from φCr30, a *Caulobacter* phage that uses RsaA as its receptor (17).

Our study defines a previously unrecognized phagehost interaction in which an N4-like phage exploits envelope polysaccharide to infect *C. crescentus*. This expands the known repertoire of *Caulobacter* phages and highlights the potential of using this phage as molecular tool to probe cell surface structure in the well-studied *C. crescentus* model system.

## Methods

### Strains and growth conditions

*Escherichia coli* was grown in Lysogeny broth (LB) or LB agar (1.5% w/v) at 37°C (53). Medium was supplemented with the following antibiotics when necessary: kanamycin 50 µg ml^-1^ or chloramphenicol 20 µg ml^-1^. *Caulobacter crescentus* was grown in peptone-yeast extract (PYE) broth (0.2% (w/v) peptone, 0.1% (w/v) yeast extract, 1 mM MgSO_4_, 0.5 mM CaCl_2_) or PYE agar (1.5% w/v) at 30°C (54). Solid medium was supplemented with kanamycin 25 µg ml^-1^ when necessary. Natural freshwater from which *Caulobacter* phage Circe was isolated was collected off the western public dock of Lake Lansing in Haslett, Michigan USA (42.755534 latitude, -84.404864 longitude).

### Plasmid and strain construction

Plasmids were cloned using standard molecular biology techniques and the primers listed in Table S4. For overexpression constructs, inserts were cloned behind the cumate-inducible promoter (P_Q5_), in pPTM057 which integrates at the xylose locus (*xylX*) and contains a cumate-inducible. For deletion constructs, inserts were constructed with overlap PCR with regions up- and downstream of the target gene and cloned into pNPTS138. Clones were confirmed by Sanger sequencing. Plasmids were transformed into *C. crescentus* via triparental mating (54). Transformants were selected by plating on PYE agar supplemented with the appropriate antibiotic and nalidixic acid to select against *E. coli* donor strains. Gene deletions were constructed via a standard two-step recombination/counterselection method using *sacB* as the counterselection marker. Transformants from the triparental mating were incubated in PYE broth without selection at 30°C for 8 hours. Cultures were plated on PYE supplemented with 3% (w/v) sucrose to select for recombinants that had lost the plasmid. Mutants were confirmed via PCR amplification of the target gene from sucrose resistant, kanamycin sensitive clones.

### Transmission electron microscopy

Phage were prepared and imaged at the Michigan State University Center for Advanced Microscopy. Briefly, 10 µl concentrated phage was incubated on a formvar carbon coated grid for 5 min, washed with distilled water, and excess water was blotted away with Whatman filter paper. Grids were stained with 1% uranyl acetate and imaged on 1400 Flash JEOL transmission electron microscope. Phage capsid diameters were measured from vertex-to-vertex in FIJI and the three vertex-to-vertex measurements for each capsid were averaged.

### Efficiency of plating assays

Strains were inoculated in PYE and incubated at 30°C overnight. Overnight cultures were diluted 1/10 in PYE and incubated at 30°C for ∼4 hours, then diluted to 0.2 OD_660_. For plating, 5 ml PYE top agar (0.3% (w/v) agar), 1 ml *C. crescentus* strain (0.2 OD_660_), and 150 µl 30% (w/v) xylose were mixed and poured on PYE plates. Phage stocks were diluted to 1 x 10^9^ PFU/ml, serially dilutions were prepared from the stock, and 5 µl dilutions were plated on PYE top agar. Plates were incubated at 30°C overnight and imaged.

### Clearing assay

Strains were inoculated in PYE supplemented with 50 µM cumate or 0.15% (w/v) xylose when applicable and incubated at 30°C overnight. Overnight cultures were diluted 1/10 in PYE supplemented with 50 µM cumate or 0.15% (w/v) xylose when applicable and incubated at 30°C for ∼3-7 hours. Cultures were diluted to 0.1 OD_660_ (∼6.7×10^7^ cells/ml) and Circe or Pasiphae were added to 0.1 multiplicity of infection (MOI) (6.7 × 10^6^ PFU/ml), 1 MOI (6.7×10^7^ cells/ml), or 10 MOI (6.7×10^8^ cells/ml). Cultures were incubated at 30°C for 24 hours while shaking and OD_660_ was measured. For clearing curves, 1 ml sample was added to three wells of 24-well plate for technical replicates. Plates were shaken (Shaking mode: orbital; <u>Orbital frequency</u>: 559 cpm (1mm); <u>Orbital speed</u>: slow) at 30°C, and OD_660_ was measured every 15 minutes for 16 hours in a Synergy HTX multi-mode plate reader (BioTek). For plate reader curves, the absorbance for three wells with PYE (i.e. blanks) was averaged and subtracted from wells that contained cells to get the blank-corrected value. Each biological replicate data point is the averaged absorbance of three blank-corrected technical replicates. Statistical analysis was carried out in GraphPad 10.6.0.

### Phage adsorption assays

Strains were inoculated in PYE and incubated at 30°C overnight. Overnight cultures were diluted 1/10 in PYE and incubated at 30°C for ∼4 hours. Cultures were diluted to 0.2 OD_660_ (∼1.3×10^8^ cells/ml), 1.3 × 10^6^ PFU/ml (0.01 MOI) Circe were added, and cultures were incubated at 30°C while shaking. At 0, 7.5, 15, 30, 45, and 60 min post-infection, samples were taken and diluted 1/100 in PYE, pelleted at 17,000 x g for 1 min, and 300 µl supernatant was mixed with 20 µl chloroform. For plating, 5 ml PYE top agar (0.3% (w/v) agar), 1 ml CB15 (0.2 OD_660_), and 150 µl 30% (w/v) xylose were mixed and poured on PYE plates. To determine the number of unadsorbed phage, samples were diluted where appropriate and 100 µl was plated on PYE top agar. Plates were incubated at 30°C overnight and plaques were enumerated. Percent unadsorbed phage was calculated as (PFU_t=x_ / PFU_t=0_)*100. Statistical analysis was carried out in GraphPad 10.6.0.

### Genomic DNA isolation

For bacterial genomic DNA, strains were inoculated in PYE and incubated at 30°C overnight. Cultures were pelleted at 12,000 x g for 1 min and supernatant was removed. Pellets were washed in 0.5 ml H_2_O, then resuspended in 100 µl TE (10 mM Tris pH 8, 0.1 mM EDTA) with 20 ng/µl RNase A. 500 µl GES solution (5.08 M guanidium thiocyanate, 0.1 M EDTA, 0.5% (v/v) sarkosyl) was added, samples were vortexed, then samples were heated at 60°C for 15 min. 250 µl 7.5 M cold ammonium acetate was added, samples were vortexed for 15 sec, and incubated on ice for 10 min. 500 µl chloroform was added, samples were vortexed for 15 sec, and pelleted at 12,000 x g for 10 min. The aqueous phase was transferred to a fresh tube, and 0.54 volumes of cold isopropanol was added. Samples were mixed by inversion and incubated at room temperature for 15 min. Samples were pelleted at 12,000 x g for 3 min and supernatant was removed. Pellets were washed with 700 µl 70% (v/v) ethanol, then air dried for 10 min. Pellets were resuspended in 100 µl TE buffer.

For phage genomic DNA, CB15 was inoculated in PYE and incubated at 30°C overnight. Cultures were diluted 1/10 in fresh PYE and inoculated at 30°C until culture reached 0.1 OD_660_. Circe was added to (0.1 MOI) and cultures were incubated at 30°C for 24 h. Cultures were pelleted at 7197 x g for 5 min and supernatant was transferred to a fresh tube. Circe was concentrated in an Amicon Ultra 15 (30K) concentrator, then treated with RNase A and Turbo DNase for 60 min at 37°C. EDTA was added to a final concentration of 15 mM, then samples were incubated at 95°C for 15 min. Lysozyme (0.1 mg/ml final concentration) and SDS (0.5% final concentration) were added and samples were incubated at 55°C for 60 min. An equal volume of chloroform was added, samples were vortexed, and samples were pelleted at 12,000 x g for 10 min. The aqueous phase was transferred to a fresh tube, and a second chloroform extraction was performed. Ammonium acetate (2.5 M final concentration) was added to aqueous phase, samples were mixed by inversion, and 0.6 volumes isopropanol was added. Samples were mixed by inversion, then stored at -20°C for 60 min. Samples were pelleted at 17,000 x g for 5 min at 4°C and supernatant was removed. Pellets were washed with 700 µl 70% (v/v) ethanol, then air dried for 10 min. Pellets were resuspended in 50 µl 10 mM Tris pH 8.5 buffer.

### Phage genome assembly and analysis

Library preparation and sequencing of genomic DNA from Circe was performed by SeqCoast. Libraries were prepared with the Illumina DNA Prep tagmentation kit and IDT For Illumina Unique Dual Indexes and sequenced on Illumina NextSeq2000 platform to get 150 bp paired-end reads. Reads were trimmed with BBDuk2 (length: >100 bp; QC: >30) and assembled using Unicycler (55) (version 0.5.1) on Galaxy (Bridging mode: normal; Lowest k-mer size: 0.2; Highest k-mer size: 0.95; k-mer steps in assembly: 10; filter out contigs lower than this fraction of chromosomal depth: 0.25). The physical ends of the chromosome were determined through Sanger sequencing. The Circe genome was annotated with a combination of Pharokka (version 1.7.3) (56), Pyrodigal (57), Phold (version 0.2.0; https://github.com/gbouras13/phold), and Phynteny (version 0.1.13; https://github.com/susiegriggo/Phynteny) using Google Colab. The viral proteomic tree was constructed using the ViPTree webserver (version 4.0) (58). Putative spanins were identified using SpaninDB (59); signal peptides were identified using SignalP (60). Sequence reads used to assemble the *Caulobacter* phage Circe genome are available through NCBI BioProject accession PRJNA1345760. The phage Circe genome is awaiting assignment of an accession number by NCBI.

### Whole genome resequencing and polymorphism identification

Library preparation and sequencing of genomic DNA from phage resistant backgrounds was performed by SeqCoast. Libraries were prepared with the Illumina DNA Prep tagmentation kit and IDT For Illumina Unique Dual Indexes and sequenced on Illumina NextSeq2000 platform to get 150 bp paired-end reads. Reads were mapped to the *C. crescentus* NA1000 reference genome (GenBank accession CP001340) and polymorphisms were identified with Breseq (61).

### RB-TnSeq to identify host genes required for Circe infection

We used a previously described and characterized randomly barcoded transposon (RB-TnSeq) mutant library in *Caulobacter crescentus* CB15 (33, 62) to identify genes important for infection by the N4-like phage Circe. For each experiment, a 1 ml aliquot of the mutant library was inoculated into 22 ml of peptone yeast extract (PYE) medium and grown shaking at 30°C overnight. For each experiment, 7 ml overnight culture was diluted into 60 ml PYE and grown at 30°C with shaking until reaching early-exponential phase (OD_660_ ∼0.1). At this point, CirceC phage was added at a multiplicity of infection (MOI) of 1.0, and cultures were returned to 30°C with aeration.

Samples were collected at 0 (uninfected), 1.5, 3, 4.5, and 7.5 hours post-infection. 10 ml of culture for the 0, 1.5, 3, and 4.5 hour samples or 20 ml of culture for the 7.5 hour samples was removed, pelleted by centrifugation (12,000 x g, 3 min, 4°C), washed once with 1 ml PYE, and stored as a cell pellet at –20°C. Genomic DNA was extracted and used for barcode amplification following the method of Wetmore et al. (63). Briefly, cell pellets were resuspended in 10–20 µl sterile water and used as template for PCR amplification of barcodes using Q5 polymerase (New England Biolabs) in 20 µl reaction volumes. Each reaction contained 1X Q5 buffer, 1X GC enhancer, 0.8 U Q5 polymerase, 0.2 mM dNTPs, 0.5 µM each of the universal forward primer (Barseq_P1) and a unique indexed reverse primer (Barseq_P2_ITxxx). Thermocycling conditions were: 98°C for 4 min; 25 cycles of 98°C for 30 s, 55°C for 30 s, and 72°C for 30 s; followed by 72°C for 5 min and hold at 4°C.

PCR products were pooled and subjected to single-end 50-bp sequencing on an Illumina NovaSeq using TruSeq primers. Barcode sequences have been deposited in the NCBI Sequence Read Archive under BioProject accession PRJNA1345765. Barcode counts were processed and analyzed using the FEBA pipeline as described in Wetmore et al. (63). Briefly, barcode counts from each post-infection sample were compared to the reference (t = 0) barcode counts to compute fitness scores (log2 ratio of barcode abundance) for each mutant. Statistical t-scores, which take into account variance in fitness of each transposon insertion strain for each gene (63), reflecting the mean fitness of a gene divided by a standard error metric. This statistic accounts for variability in fitness measurements across mutant strains and provides some indication of how many standard-error units the fitness effect differs from zero.

## Host infection fitness time-course clustering

Gene fitness scores (*t*-scores) were obtained at 1.5, 3.0, 4.5, and 7.5 hours following N4-like phage (CirceC) infection. Genes with |*t*| ≥ 4 at one or more time points were retained for analysis. To compress dynamic range while preserving magnitude and sign, a sign-preserving logarithmic transformation was applied. Hierarchical clustering of genes (rows) was performed on the transformed matrix using Euclidean distance and average linkage through SciPy. Gene order was determined by dendrogram leaf order. Time points (columns) were not clustered and are displayed chronologically. Complete RB-TnSeq fitness scores and corresponding t-scores are presented in Table S3.

## Supporting information

CirceC Draft Genbank Flat File

## Acknowledgements

Microscopy data collection was performed at the MSU Center for Advanced Microscopy by Alicia Withrow. We want to acknowledge Alicia Withrow for her helpful conversations around TEM imaging. Funding for work came from University of Michigan–Flint startup funds to M.M., NIH award R35GM131762 to S.C., and by Army Research contract W911NF2210105 to S.C. This study is dedicated to Lucia Rothman-Denes, who introduced A.F. and S.C. to phage N4.

---

**Figure S1.**
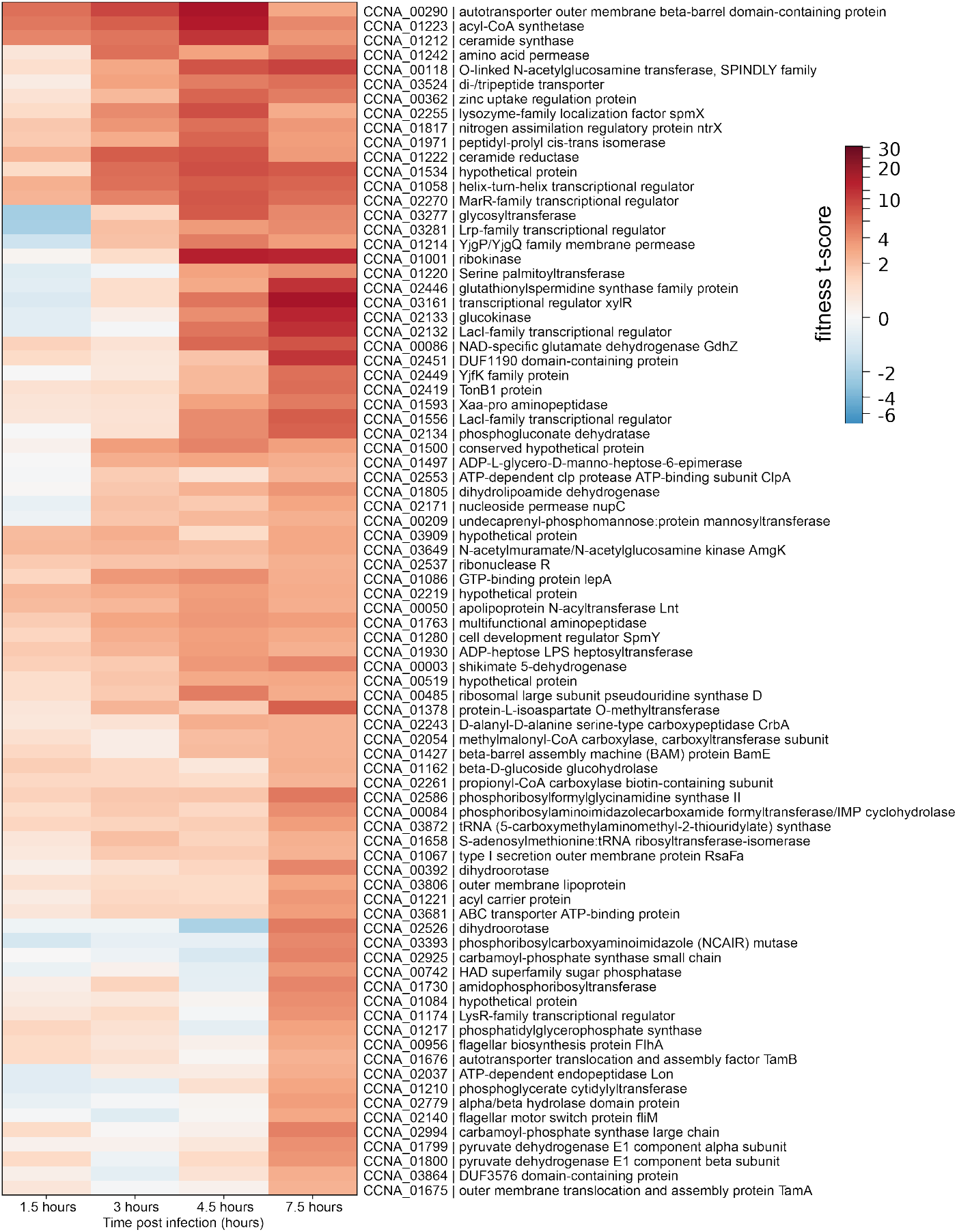
Hierarchically clustered heatmap of *C. crescentus* gene fitness t-scores at 1.5, 3.0, 4.5, and 7.5 h post-infection with CirceC at MOI=1. Rows include genes with ∣ *t* ∣≥ 4at ≥1 time point. Figure shows clustered genes not presented in panel 7B. Complete fitness data are presented in Table S3.

**Figure S2.**
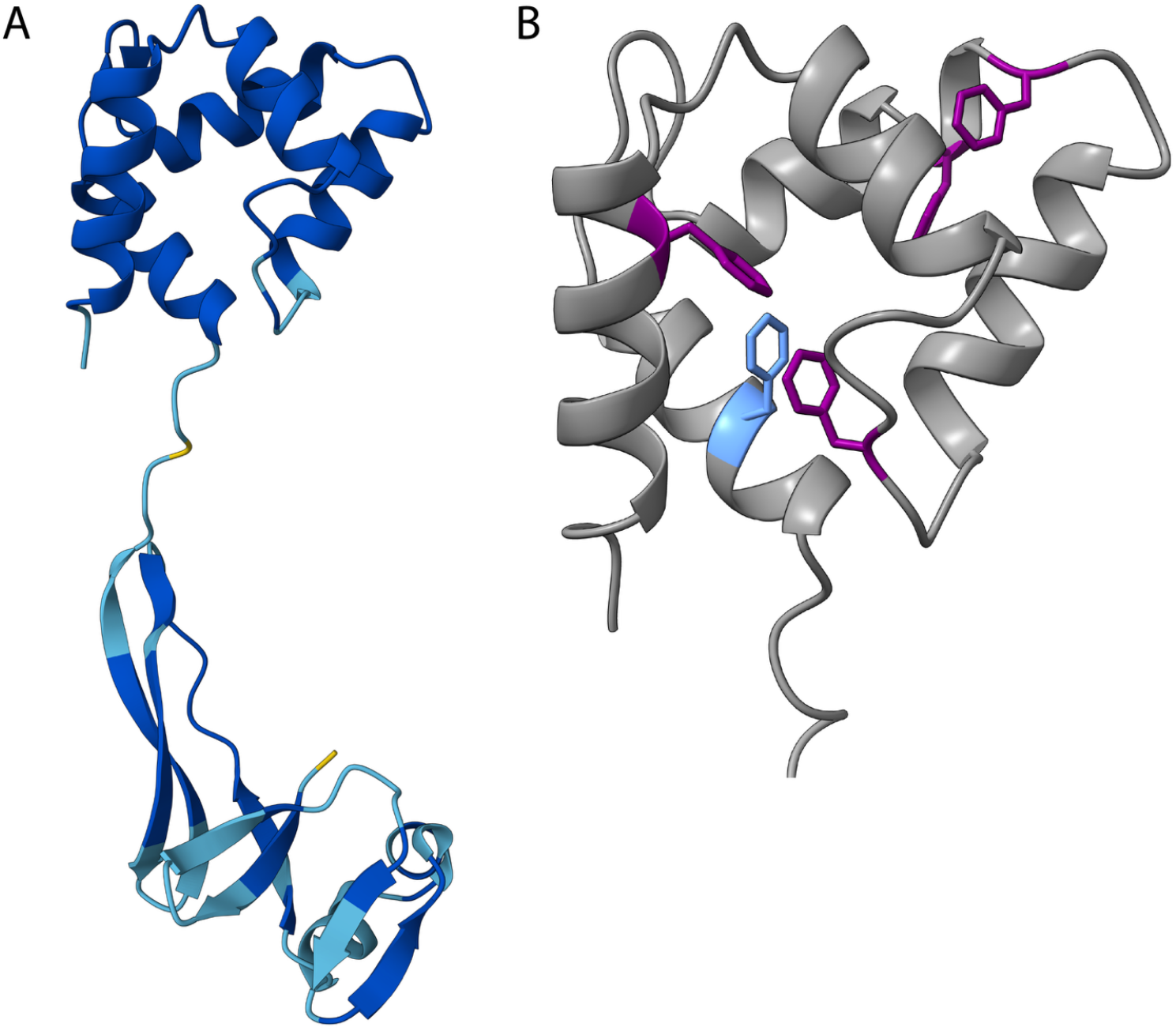
Predicted gp063 structure. Structure of gp063 predicted by AlphaFold3. A) Full-length gp063 colored based on pIDDT score (pTM=0.53). B) C-terminal domain of gp063. Phenylalanine residues are colored purple. Phenylalanine 91 is colored blue.

